# A modified transplant design reveals that habitat quality and quantity both limit a species’ range

**DOI:** 10.64898/2026.04.22.720096

**Authors:** Olivia Rahn, Anna L. Hargreaves

## Abstract

Species’ range edges provide excellent arenas for testing which ecological constraints prevent expansion into new environments. Theory predicts that ranges can be constrained by declines in the quality or amount of habitat, but their relative impact is unknown because empirical studies are seldom designed to quantify habitat amount. Here, we propose a simple modification of across-the-range-edge transplant experiments that enables tests for declines in both habitat quality and amount. Using this design, we show that quality and amount of suitable microhabitat both decline across the high-elevation range edge of the herb *Rhinanthus minor*. Using simulation models parameterized with field data, we show that either decline is sufficient to impose range limits, and both declines contribute to limiting *R. minor*’s high-elevation range. We end with three simple suggestions for the design and presentation of across-the-range-edge transplant experiments that would clarify how often and severely declines in habitat amount limit species’ ranges.

## Introduction

A foundational goal of ecology is to identify which factors determine whether species can expand into new habitat. This goal is central to explaining the size and location of species’ geographic ranges, particularly the location of their range edges, and is increasingly important as ecologists try to predict range expansions of introduced species and range shifts of native species responding to climate change (Chen et al., 2011; Davis & Shaw, 2001; Early & Sax, 2014).

When a species reaches a stable range limit without any clear barrier to dispersal, the primary ecological explanations can be roughly divided into declines in the quality of habitat (Brown, 1984) or in the amount of habitat (Carter & Prince, 1981; Holt & Keitt, 2000). Models predict that either type of gradient can limit a species’ range (e.g. Lennon et al., 1997), and it is often assumed that habitat quality and amount both decline toward and beyond species’ range edges (Hampe & Petit, 2005), though their relative role in limiting ranges is rarely known.

Theory that ranges are limited by habitat quality stems from the long-standing idea that a species’ range is a spatial representation of its (realized) niche (McArthur 1972; Humbolt & Bonplant, 1805). In this view, species expand their ranges across environmental gradients until one or more ecological factors exceed the species’ tolerance. Stable range limits are expected to arise where habitat quality becomes too low to support self-sustaining populations (Holt et al., 2003; Soberón, 2007). The strongest test of this prediction is to transplant species into habitat beyond their range and compare performance to transplants at the range core and edge. Syntheses of such transplant experiments show that habitat quality often declines beyond species’ ranges (Hargreaves et al., 2014; Lee-Yaw et al., 2016) caused by diverse abiotic and biotic factors (Paquette & Hargreaves, 2021). Yet these experiments also suggest that for ∼25% of range limits, beyond-range habitat is of sufficient quality to support viable populations. Thus, while declining habitat quality contributes to many (perhaps most) range limits, other processes are also undeniably important.

For many species, the amount of habitat also appears to decline toward and beyond their range edges via increasing habitat patchiness at either the landscape or micro-habitat scales (Griggs, 1914, Rapoport 1982, Gillies et al., 2025). Declines in habitat amount can limit ranges by impeding colonization via increasing patch isolation, or by increasing stochastic extinctions if patches become smaller and so support fewer individuals (Carter & Prince, 1981; Holt & Keitt, 2000; Oldfather et al., 2021), even when habitat quality within the best-available patches remains high at and beyond the range edge. Despite the recognized potential importance of habitat amount, and calls for more experiments that consider its role in limiting species ranges (e.g. Pironon et al., 2017), empirical tests generally focus on habitat quality, leaving the importance of habitat amount unclear (Fig. 1).

**Figure 1.**
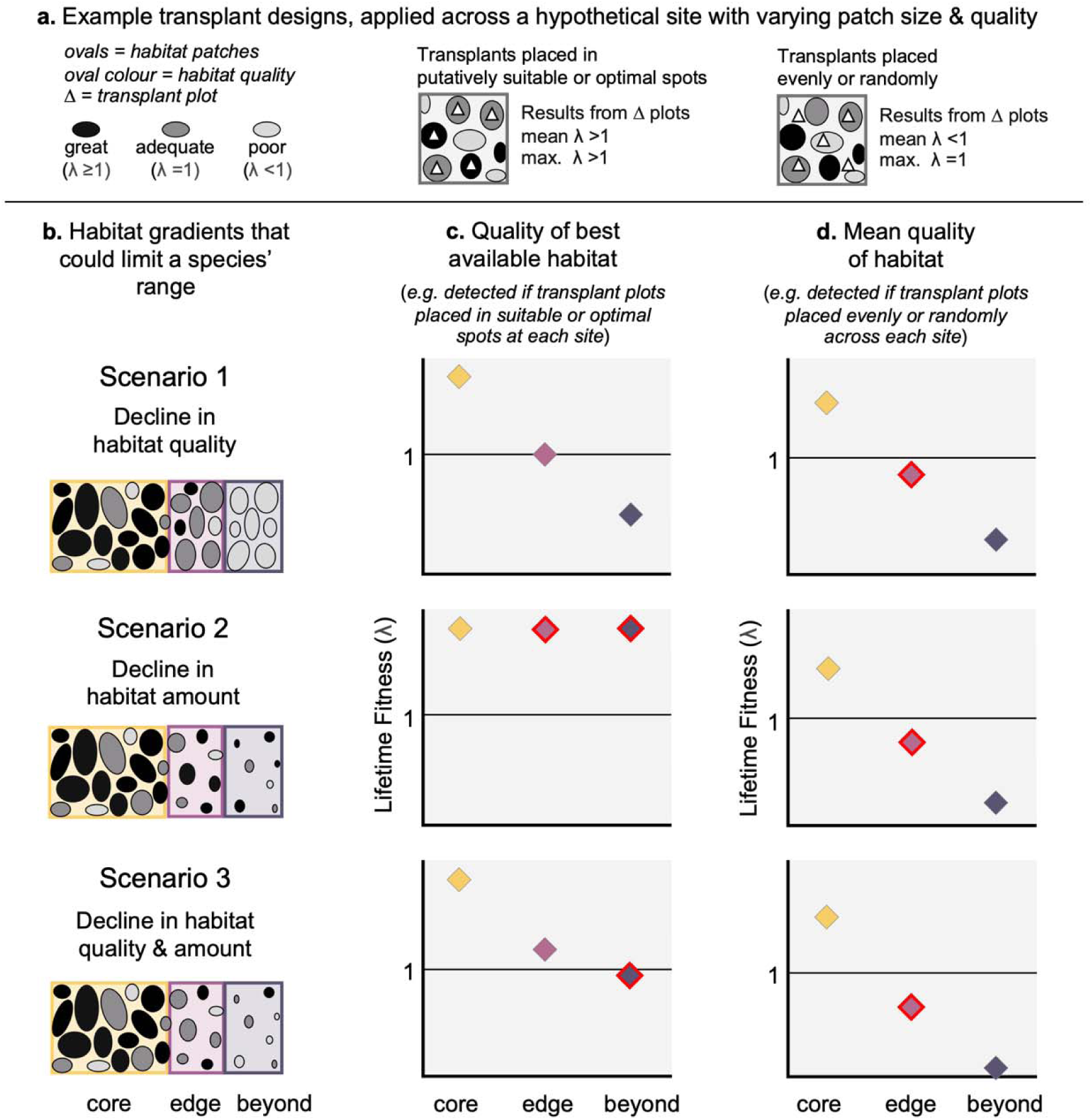
Illustrations of how transplant designs vary in the habitat gradients they detect. **(a)** Schematics of two common designs applied to a cartoon site, showing the contrasting conclusions they would reach about fitness (λ). For simplicity, assume λ between patches is 0. **(b)** Three scenarios (*Sc*) that vary in whether a species’ range is limited by declines in habitat quality, habitat amount, or both across a landscape. (**c–d**) Example results from each transplant design applied across each landscape, with one site per range position. Red point outlines indicate where results from a particular design may lead to misinterpretations about the range-limiting gradients. (**c**) Transplant plots placed in putatively suitable/optimal habitat capture declines in the quality of the best available habitat *(Sc1&3)*, and are most likely to identify suitable habitat beyond the range *(Sc2)*, but may miss declines in habitat amount (e.g. *Sc2*: core, edge, and beyond habitat appear equally suitable) and may overestimate overall site suitability (e.g. *Sc2&3:* unsuitable beyond-range habitat appears suitable). (**d)** Transplant plots placed evenly or randomly across sites capture the combined effect of habitat quality and amount, but may miss rare suitable patches and underestimate overall site suitability (e.g. *Sc2&3*: edge habitat supports self-sustaining populations but appears to be unsuitable with λ<1).

This knowledge gap stems partly from how across-the-range-edge transplant experiments are typically designed (Fig. 1a). To avoid the critique that beyond-range transplants failed because they did not sample suitable habitat (rather than because suitable habitat does not exist), researchers often transplant individuals into putatively suitable or optimal sites/microsites, (e.g. Lajoie & Vellend, 2018; McLane & Aitken, 2012; Samis & Eckert, 2009). This design provides the strongest test of whether suitable habitat exists beyond species’ ranges, and whether the quality of the best available habitat declines toward and beyond range limits. However, it may miss declines in the *amount* of suitable habitat present, and so can overestimate the average suitability of a site or region (Fig. 1c). This could lead researchers to misinterpret a stable range limit driven by declining habitat amount (i.e. beyond-range habitat is too patchy to be suitable) as a temporary range limit in the process of expansion (i.e. beyond-range habitat is suitable for eventual colonization). An alternative design is to place transplants evenly or randomly across a transplant site (*e.g.* Davison & Field, 2018; Levin & Schmidt, 1985; Solarik et al., 2018). This design should detect the combined effects of habitat quality and amount, but may miss rare suitable patches and underestimate whether a site or region can support self-sustaining populations (Fig. 1d).

We propose a simple design modification that would enable transplant experiments to disentangle the effects of habitat quality and amount. We focus on the microsite level, where the design is easily implemented without significant additional effort. Rather than placing all transplant plots in putatively suitable microsites or evenly/randomly, we suggest deliberately placing some plots in putatively suitable (or optimal) microsites and distributing the others evenly or randomly across each site (within the species’ normal habitat, but agnostic to microsite quality). Putatively suitable microsites can be identified however researchers normally would. e.g. via favourable biotic or abiotic conditions, or the presence of the focal species at in-range sites. Incorporating putatively-suitable plots provides the strongest test of whether suitable microsites exist beyond the range and whether the quality of the best available microsites declines across the range edge. Evenly or randomly distributed plots test whether the amount of suitable microhabitat varies among sites. By explicitly disentangling gradients in habitat quality and amount, this design improves explanatory power and reduces risk of misinterpreting transplant results (Fig. 1).

We implemented this design to test whether (micro)habitat quality or amount decline across the high-elevation range limit of the annual plant *Rhinanthus minor. Rhinanthus* species are hemiparasites, extracting nutrients and water from neighbouring plant roots. *Rhinanthus minor* uses a wide host range of >50 species, but optimal hosts include grasses, legumes and yarrow (*Achillea millefolium)* (Gibson & Watkinson, 1989; Saona, 2002). Previous transplant experiments, which used the presence of suitable hosts to identify putatively-suitable microhabitat (Hargreaves & Eckert, 2019; Persi, 2020), showed that the quality of the best available habitat declines dramatically above *R. minor*’s range. However, along the best-studied elevational transect (Mt. Allan; Fig. 2), several results suggest that declining habitat quality cannot entirely explain *R. minor*’s high-elevation range limit: suitable hosts exist above *R. minor*’s range but become more patchily distributed (personal observations), and in some years suitable microhabitat (i.e. transplant plots with lifetime fitness ≥1) existed above *R. minor*’s range (Hargreaves & Eckert, 2019). We therefore suspect that declining habitat amount also plays a role, but this has not yet been quantified.

**Figure 2.**
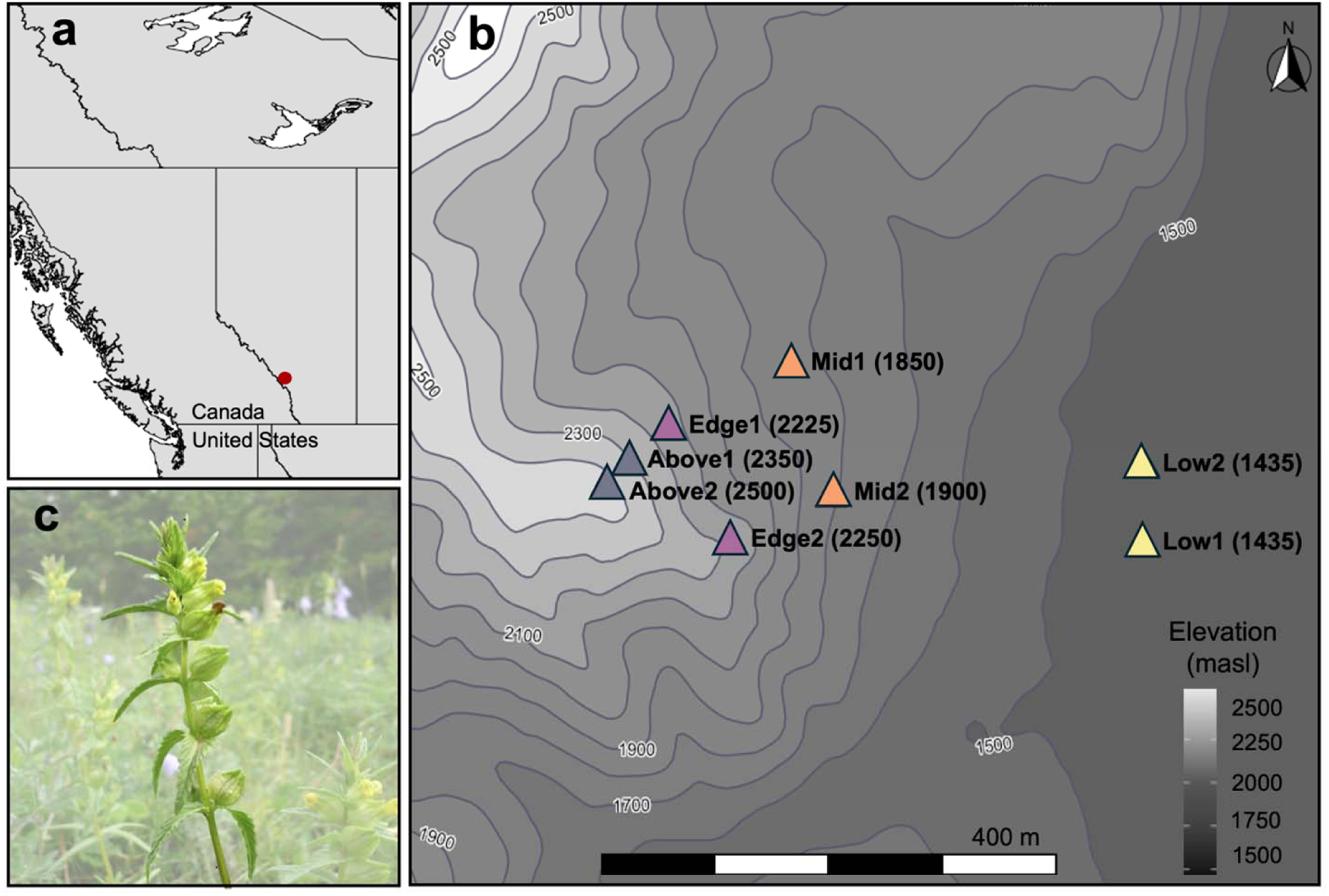
Study area and species. **(a)** Location of our study region (red dot) within northwestern North America. **(b)** Transplant sites along Mt. Allan: Low = toward but not at *Rhinanthus minor*’s lower range edge, Mid = *R. minor*’s elevational range centre, Edge = upper range edge, Above = above *R. minor*’s range. **(c)** Flowering *R. minor* plant (background faded out for clarity).

Our study has two parts. First, we use the modified transplant design to assess habitat suitability at 8 sites across and above *R. minor’s* range, placing some plots in putatively high-quality microsites and remaining plots evenly across each site. We recorded mean lifetime fitness (which approximates λ for *R. minor*) in each plot, which we used to quantify habitat quality (λ) and the proportion of plots that were suitable (λ≥1). Second, we evaluated the relative importance of declining habitat quality and amount in limiting *R. minor*’s range using a simple range-expansion model parameterized with our field data. We tested five predictions. Given previous findings that quality of putatively-suitable microsites declines above *R. minor*’s range, we predicted that: **P1)** overall site quality (mean λ across all transplant plots, which incorporates microhabitat quality and amount) will decline from in-range to beyond-range sites and be <1 (i.e. unsuitable) beyond the range; and **P2)** the best available habitat will be lower quality beyond the range than within. If declining habitat amount also contributes to *R. minor*’s range limit, we predict that: **P3)** the proportion of suitable plots will decline beyond the range; **P4)** evenly-spaced plots will detect stronger across-the-edge fitness declines than putatively-suitable plots; and **P5)** simulations that include variation in habitat amount will exhibit slower expansion and yield a predicted range edge that better matches *R. minor*’s actual range edge.

## Methods

### Study system

*Rhinanthus minor* is an annual herb native to lightly-disturbed open meadows in temperate North America and Europe (Hulst et al., 1986; Westbury, 2004). In our study region (Kananaskis, Canada), *R. minor*’s elevational range limits roughly correspond to lower treeline (∼1200 masl) and upper treeline (∼2300 masl) (Ensing et al., 2021; Hargreaves, 2014). It is primarily self-fertilizing and can produce a full set of viable seeds autonomously (Hargreaves et al., 2015), so reproduction of beyond-range transplants is not limited by pollination. We conducted transplant experiments along the eastern and southeastern slopes of Mt. Allan, which have natural and cleared meadows from 1400 to 2500 masl (Fig. 2).

### Transplant Design

During fall 2021, we planted *R. minor* seeds at 8 sites at 4 range positions: 2 sites toward (but not at) *R. minor*’s low-elevation range edge (Low1&2); two at its range centre (Mid1&2); two at its high-elevation range edge (Edge1&2), and two above its range (Above1&2; Fig. 2a). Five of these sites (Low1, Mid1, Edge1, Above1, Above2) formed the ‘Nakiska’ transect of Hargreaves & Eckert 2019. Within-range sites are dominated by grasses with many legume species also present. Above1 (2300 masl) has extensive grassy areas with stunted trees and rocky outcrops, whereas Above2 (2500 masl) is dominated by alpine heathers and avens.

At each site we planted *R. minor* seeds into 30 plots. Five plots were selected *a priori* as putatively-suitable based on the presence of *R. minor* and/or suitable hosts: graminoids, legumes or yarrow. The remaining 25 plots were spaced in an even grid across the site, though at Above1 some plots were placed off-grid to avoid rocky outcrops. Each plot was 1.2 m^2^, separated from other plots by ≥5 m. We removed natural *R. minor* plants before they dropped seed, but left other vegetation intact, so transplants capture all biotic and abiotic influences on fitness. Seeds were planted 20 cm apart in a grid just below the ground surface, marked with a toothpick. Each plot had 25 seeds; 20 from the local population (using Edge1&2 seeds for Above1&2 sites, respectively), and 5 seeds from a standard seed source (Mid1) test for effects of local seed sources due to poor provisioning or genetic quality. Local and standard seeds detected the same patterns of site suitability, suggesting results from local seeds primarily reflect site quality rather than source quality (Table S1).

### Transplant Monitoring

We monitored transplant performance during summer 2022. We checked sites every 2–3 days during emergence (late May to late June), then approximately once a week until mid-September to track survival and reproduction. For each plant that produced fruits, we estimated female reproductive success by counting the viable seeds (seed viability can be reliably assessed from colour and plumpness) in ∼25% of fruits per plant. We estimated seed production per reproductive plant as *mean viable seeds/fruit x number of fruits*. We monitored temperature at 10–20 plots per site using iButton temperature loggers (see Fig. S1).

### Estimating plot-level λ

As *R. minor* is an annual with little seedbank, the population growth rate of each plot (λ) is roughly equivalent to mean lifetime fitness or the *seeds produced per seed planted* (Hargreaves & Eckert, 2019). We used this to quantify each plots’ habitat quality (n = 30 plots/site except for Mid1, where five plots were destroyed by a mower before seeds were counted). Since all plots had the same number of seeds planted, we modeled *seeds produced per plot* as the response because count data enable a wider range of statistical approaches, but present results as λ *=* seeds produced/seed planted to facilitate comparison with other studies.

### Statistical Analyses

Analyses were conducted in R v. 4.5.2 (R Core Team, 2025). We tested whether habitat quality and amount differed among sites or range positions using generalized linear models (GLMs; glmmTMB and MASS packages; Brooks *et al.,* 2017, (Venables & Ripley, 2002)). Some analyses used range position (Low, Mid, Edge, Above) instead of site because three sites with zero suitable (λ≥1) plots interfered with model convergence. If count data were zero-inflated we added a zero-inflation component to models (Ludecke et al., 2021). We assessed significance of predictors using Type II ANOVAs for negative binomial GLMs, Type II Chi-Squared for binomial GLMs (car package, Fox & Weisburg 2019), and likelihood ratio tests for models with interactions (lmtest package, Zeileis & Hothorn 2002). When a predictor was significant, we tested which levels differed using post-hoc pairwise comparisons with a Tukey adjustment for multiple comparisons (emmeans package, Lenth, 2025).

#### Overall site quality

We tested whether overall site quality (i.e. the combined effect of microhabitat quality and amount) differed among sites **(P1)** using λ from all transplant plots: *seeds produced per plot ∼ site* (negative binomial GLM, log-link).

#### Quality of best-available habitat

Explicitly incorporating putatively-suitable plots increases the chance of capturing the best-available habitat at each site, but evenly-spaced plots may also capture high-quality patches. We therefore assessed whether the quality of the best-available habitat declined across *R. minor*’s range edge **(P2)**, using λ from plots in which transplants performed well, regardless of whether they were originally selected as putatively-suitable (negative binomial GLMs). We used two definitions of ‘best-available’: (1) plots with λ≥1 (i.e. suitable plots; including only sites with some λ≥1plots), and (2) plots in which transplants produced fruits even if final λ was <1. We included the second definition as some plants matured fruits but were browsed by mammals before seeds could be counted, therefore reducing λ estimates and sample sizes; since browsing effects do not vary systematically among sites across years (Falk 2013), we consider it a haphazard influence on fitness, rather than a range-limiting component of the environment.

#### Habitat amount

We tested whether the proportion of suitable plots (defined as those with λ≥1) declined above *R. minor*’s range (**P3**) using binomial GLMs (logit link). As three sites had no λ≥1 plots, we tested whether the proportion of suitable plots varied a) among range positions, including data from all sites (*plot quality ∼ range position*) or b) among sites, including only the 5 sites that had at least one suitable plot (*plot quality* ∼ *site*). We ran both models using only the 25 evenly-spaced plots/site (the ideal approach), and again including the 5 putatively-suitable plots/site. Trends were similar, but including all 30 plots improved sample size and our ability to detect significant pairwise comparisons. We therefore present data from all 30 plots (Fig. S2 presents results from evenly-spaced plots only).

#### Experimental design

We tested whether evenly-spaced plots detected steeper fitness declines than putatively-suitable plots (**P4**) using a negative binomial GLM: *seeds produced per plot ∼ site x plot type*.

#### Breakdown by life-history stage

While lifetime fitness is the strongest test of habitat quality, we also analyzed fitness components separately to identify which most influenced trends. We tested whether each fitness component varied among sites, using one model per component: emergence (emerged seedlings per plot; negative binomial GLM); survival (proportion of seedlings that survived to make fruits; binomial GLM); and reproduction (seeds produced per seed-producing plant; negative binomial GLM).

### Simulation Models

We tested whether including variation in habitat amount during simulated range expansion resulted in a) slower expansion or b) a more accurately predicted range edge for *R. minor* **(P5)**. We simulated expansion across three landscape types that had gradients in habitat quality, habitat amount, or both, parameterized with values from our transplant experiment. Each landscape was 100 cells wide and 500 rows long, divided into five 100-row sections corresponding to the Low, Mid, Edge, Above1 and Above2 range sections from our transplant experiment (Fig. S3). We separated Above1 and Above2 as they differ in elevation, *R. minor* performance, and habitat.

#### Varying habitat amount

Within each 100-row section, a fixed proportion of cells (randomly-selected) was designated as suitable. In landscapes with a gradient in habitat amount this proportion varied among sections, taken from the observed proportion of suitable plots in that range section during our experiment. For landscapes without a gradient in habitat amount, all sections used the ‘Low’ section proportion (Low sites together had the highest overall habitat amount and quality).

#### Varying habitat quality

Each cell in the landscape was assigned a habitat quality value λ. In landscapes with a gradient in quality, we drew λ values for suitable cells from negative binomial distributions parameterized using field-observed λ’s from suitable (i.e. λ≥1) plots in the corresponding range section. For landscapes with no quality gradient, all sections used the ‘Low’ distribution. Unsuitable cells always drew λ from a distribution parameterized using unsuitable plots from the Low section, such that matrix quality was constant across the range. After cell λs were assigned, we scaled them to vary from 0–1.

Detailed explanations of simulation methods are in Supplementary Methods 1. In brief, our model is based on previous range expansion models (Hargreaves et al., 2015; Phillips, 2012), but considers a single genotype that does not evolve. Model parameters were chosen to reflect *R. minor* ecology. Each simulation began with 1000 plants placed randomly throughout the ‘Low’ section. Individuals established and reproduced, with offspring number drawn from a Poisson distribution whose mean was calculated using a modified Hassel-Commins population-growth equation that incorporates density dependence: λ *= R_max_/(1 + N/K (R_max_ −1)).* Carrying capacity (*K)* and maximum offspring per individual (*R_max_*) were scaled based on cell quality. To reflect random mortality, half of each plant’s seeds were selected to survive, then dispersed in random directions by a distance drawn from a Poisson distribution (mean=1 cell). After 1200 generations we recorded the furthest row colonized, total population size across the landscape, and average cell density. We ran 40 replicates of each landscape type.

We also used a variation of our simulation model to test how transplant design influences interpretations about the nature of range limits. To do this, we compared range expansion across landscapes parameterized with values from a) only putatively-suitable plots (representing results from design in Fig. 1c) or b) all transplant plots (representing our modified design). Details in Supplementary Methods 2.

## Results

### Overall site quality

Mean λ across all plots (putatively-suitable + evenly-spaced plots) varied significantly among sites (χ^2^_df=7_ = 53.5, *P* < 0.0001; Fig. 3a). Most sites in the range core outperformed most sites at the range edge or beyond, and sites above *R. minor*’s range were unsuitable (mean λ<1), supporting **P1**. Although all in-range sites had natural *R. minor* populations, one mid-range and both range-edge sites also had λ<1, so may be unsuitable over longer terms.

**Figure 3.**
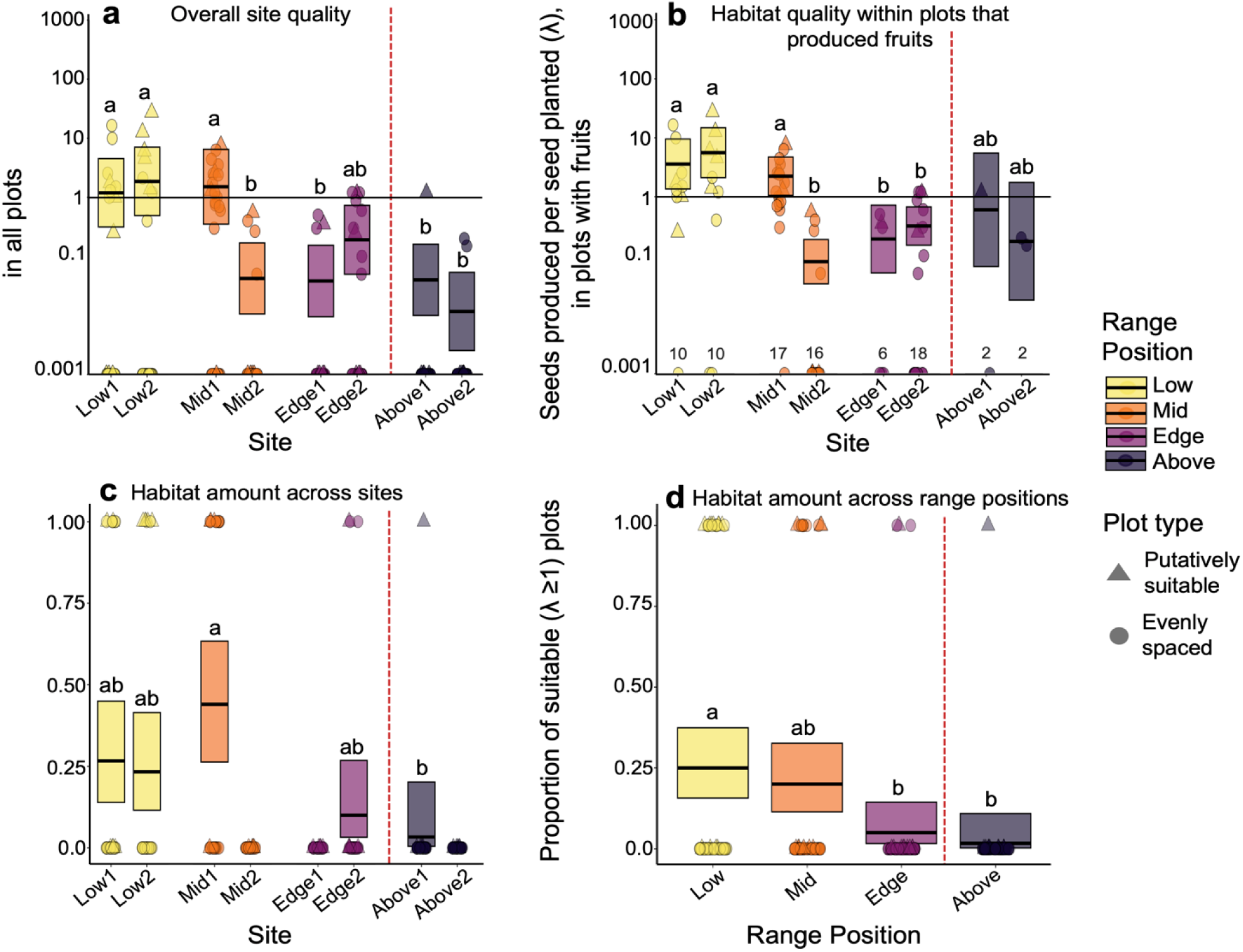
Habitat quality and amount both decline across *R. minor*’s high-elevation range edge. Sites are from two elevational transects spanning most of *R. minor*’s elevational range, with the red line separating within-range (left) and beyond-range (right) sites. **a)** Overall site quality (mean λ across transplant plots placed in microhabitats deemed *a priori* as putatively suitable and plots placed evenly across each site agnostic to microhabitat quality). Sites with overall λ <1 are predicted to be unsuitable. **b)** Quality of best-available microhabitats (λ in plots where transplants made fruit; Fig. S4 gives results for alternate definition of best-available). **(c­d)** Habitat amount. The amount of suitable microhabitat varied among sites (**c**) and range positions (**d**). In c, three sites with no suitable plots were excluded from the model to improve model convergence. Mid-lines and shaded rectangles show marginal means and 95% confidence intervals back-transformed from GLMs; points show raw data from each plot. In a,c,d, n=30 plots/site except Mid1 where n=25 due to accidental mowing before seeds could be counted; in b numbers show n=plots with fruits/site.

### Quality of best-available habitat

The quality of suitable microsites varied among sites, but the strength of this pattern depended on the definition of ‘best-available’ and resulting sample size (mixed support for **P2**). If we defined ‘best-available’ as plots with λ≥1, quality (λ) declined slightly toward and beyond *R. minor*’s range edge (*site*: χ^2^_df=4_ = 12.9, *P* = 0.012), but most pairwise comparisons were non-significant (the exception was Low2 outperforming Edge2; Fig. S4). If we defined ‘best-available’ as plots in which transplants produced fruits, habitat quality declined more strongly from the range core to edge (*site*: χ^2^ = 71.0, *P* < 0.0001; Fig. 3b). In both analyses, quality of above-range sites was not statistically lower than 1 nor less than other sites (Fig. 3b).

### Habitat amount

The amount of suitable microhabitat declined across *R. minor*’s range edge (supporting **P3)**. The proportion of suitable (λ≥1) plots differed among range positions (χ^2^_df=7_ = 14.4, *P* = 0.0025; Fig. 3d) and among sites that had at least 1 suitable plot (χ^2^ = 12.8, *P* = 0.012; Fig. 3c). There was less suitable habitat above the range than in the range core, although sites without suitable plots also existed in the range-centre (Mid2).

### Effect of transplant design

Evenly-spaced plots detected lower habitat quality and qualitatively steeper fitness declines across the range edge than plots placed in putatively-suitable microhabitats (Fig. S5; *plot type*: χ^2^_df=1_ = 5.94, p = 0.015), partially supporting **P4** and illustrating the importance of transplant design (Fig. 1). Evenly-spaced plots predicted that sites above *R. minor*’s range were unsuitable (mean λ significantly <1), whereas putatively-suitable plots did so with less certainty (Fig. S5). In simulation models used to explore the effects of transplant design (Supplementary Methods 2), landscapes parameterized using transplant fitness from putatively-suitable plots only (i.e. mimicking a typical design that targets the best-available habitat) supported faster range expansion and higher population densities than those parameterized using data from all transplant plots (Fig. S6), but both underestimated *R. minor’s* real range limit.

### Breakdown by life-history stage

Elevational patterns in performance varied among *R. minor* life stages (Fig. 4). Sites differed in emergence (χ^2^_df=7_ = 182.1, *P* < 0.0001), survival (χ^2^_df=7_ = 29.4, *P* = 0.0001), and reproductive success (χ^2^_df=7_ = 53.5, *P* < 0.0001). Emergence showed the strongest spatial patterns, and was consistently higher at in-range sites than above the range (Fig. 4). Above-range sites also had among the lowest survival and reproduction, though seed production was also very low at a mid­range and range-edge site.

**Figure 4.**
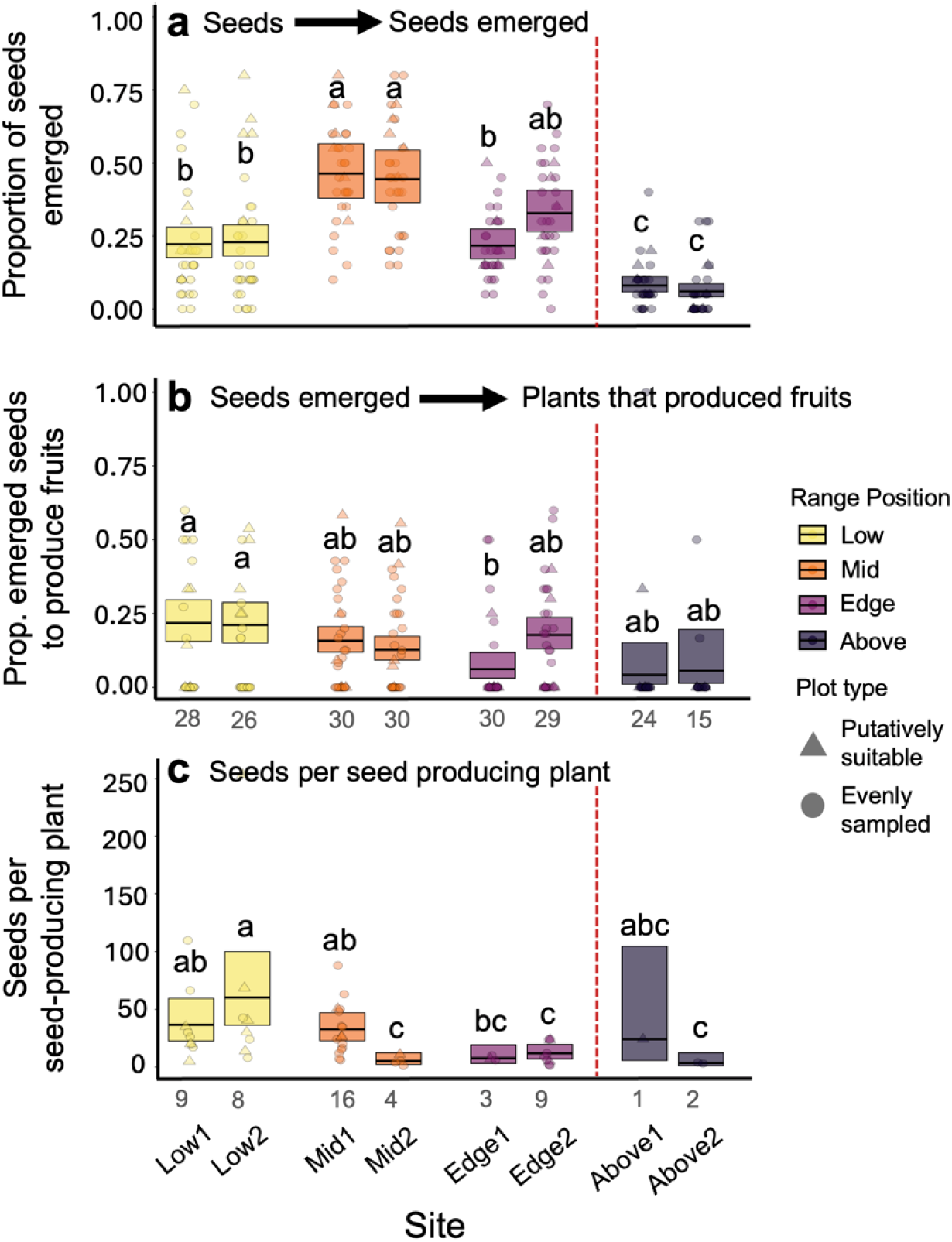
Performance of *R. minor* transplants by life stage. Data include both evenly-spaced and putatively-suitable plots at each site. **a)** Emergence: proportion of planted seeds that emerged to produce seedlings (n=20 seeds/plot and 30 plots/site). **b)** Survival to reproduction: proportion of emerged seedlings to produce fruits (n=plots with emerged seedlings, indicated below the x-axis). **c**) Reproduction: mean seeds produced per seed-producing plant (n=plots with plants that produced fruits). Formatting as in Fig. 3.

### Simulation Model

We predicted that if habitat amount declines across *R. minor*’s range edge, simulations that include gradients in habitat amount will exhibit slower expansion and a predicted range edge that better matches *R. minor*’s actual range edge (**P5**). We found some support for this. Differences among landscape types were strongest before stable range limits formed (Fig. 5), though less apparent by generation 1200 (Fig. S4). Both gradients were strong enough to limit *R. minor*’s range independently. Expansion was slightly faster and more extensive across gradients in only habitat amount (mean ± SD furthest row reached by generation 1200: 215.4 ± 6.9, Fig. 5) compared to gradients in only habitat quality (206.8 ± 1.8), and was slowest in landscapes with gradients in both quality and amount (202.9 ± 1.0; Fig. 5, Table S2). Landscapes with gradients in both habitat quality and amount also had the lowest final population sizes, particularly at the range edge (Fig. S7, Table S2). All scenarios underestimated *R. minor’s* ‘real’ range limit, reaching range limits between rows 202 and 215 whereas the boundary between in-range and above-range conditions occurred at row 300. These results do align with the low transplant fitness at edge sites in 2022 (Fig. 3), and recent declines in the size and performance of natural range-edge populations of *R. minor* (see Discussion).

**Figure 5.**
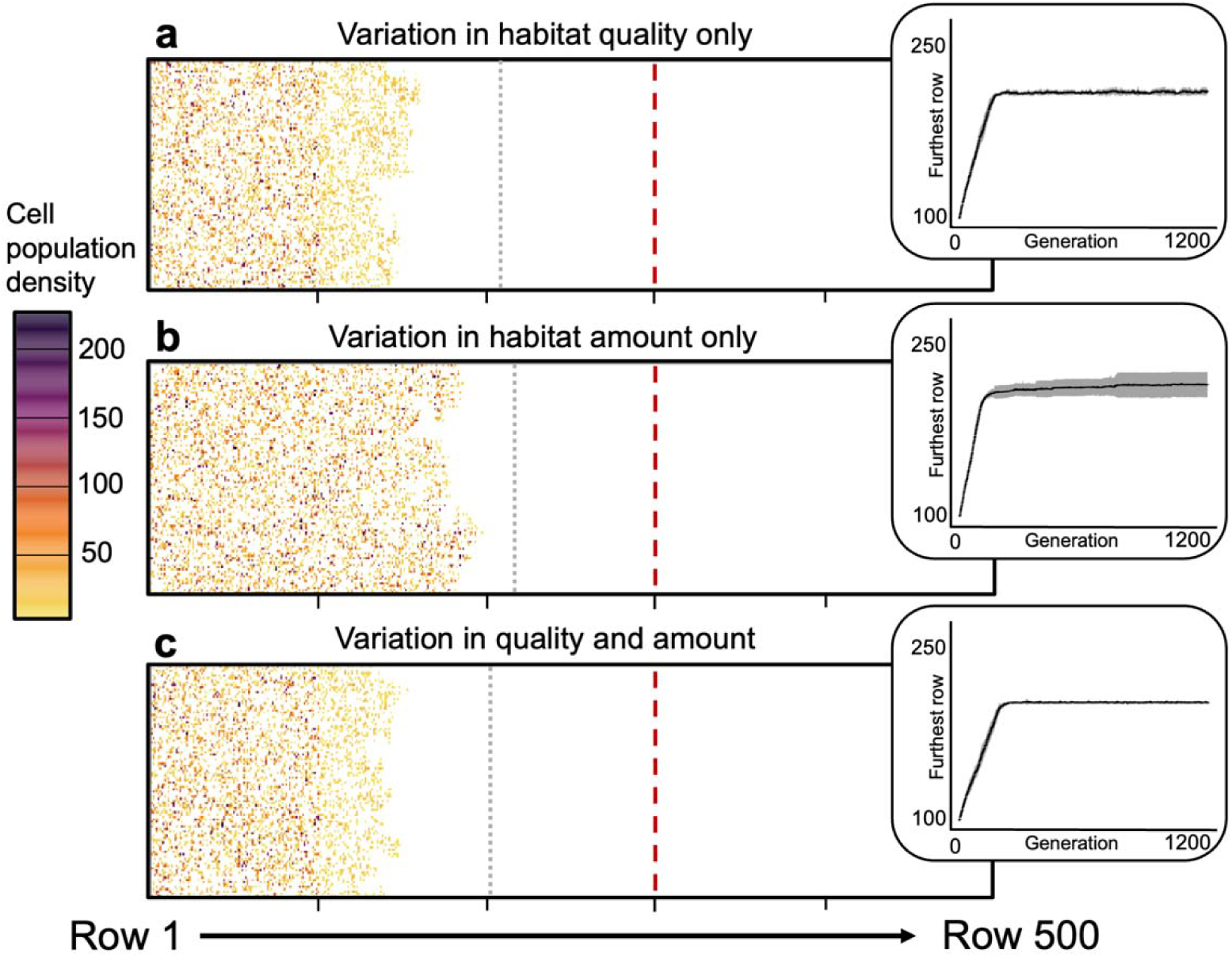
Simulated *R. minor* range expansion across landscapes parameterized with field data. Landscapes are divided into range sections of 100 rows each (Low, Mid, Edge, Above1, Above2; delineated by x-axis ticks). Landscape types differ in whether range sections vary in the quality (**a**), amount (**b**), or quality and amount (**c**) of suitable cells. Landscape plots show densities in all cells at generation 100 for a randomly selected model run, as this is when differences among landscapes were most apparent. Dashed red lines show *R. minor*’s ‘true’ range limit at row 300, dotted grey lines show the average extent of range expansion for that landscape type at generation 1200. Insets: accumulation curves showing the furthest row of the landscape reached at each generation (mean ± 2 standard deviations across 10 randomly selected runs per landscape type).

## Discussion

### Transplant design and habitat amount

Declining habitat amount has long been posited as an important driver of species’ range limits (Carter & Prince 1981; Holt & Keitt 2000). We argue conceptually that quantifying habitat amount can alter interpretations about species’ range limits (Fig. 1), and demonstrate this empirically using the high-elevation range limit of *Rhinahthus minor*. We disentangle habitat amount and quality and show that both factors decline with increasing elevation and contribute to limiting *R. minor*’s range (Fig. S3,5). A transplant experiment that targeted only putatively-suitable microsites might not have ruled out the possibility that above-range habitat could support self-sustaining populations (Fig. S5). It also may have detected shallower declines in overall site suitability by missing significant declines in the amount of suitable microhabitat (Fig. 3c,d). Indeed, our simulation models showed that declines in microhabitat amount were strong enough to limit *R. minor*’s range alone (Fig. 4), even without the strong co-occurring declines in habitat quality (Fig. 3b). Our results provide rare empirical evidence that declines in microhabitat amount contribute to a species’ range limit, and highlight the importance of incorporating habitat amount in empirical studies. Below, we discuss important considerations in measuring habitat amount; an unexpected implication of our results for *R. minor*’s range dynamics; and simple suggestions to enable future experiments, regardless of design, to better reveal the role of habitat amount in limiting species’ ranges.

One challenge in quantifying habitat amount is that patchiness is highly scale dependent (Lechowicz & Bell, 1991; Wiens, 1976). Theory about habitat amount and range limits comes mostly from the metapopulation literature, which focuses on patchiness at the landscape scale, e.g. patches of meadow in forest matrix (Hanski, 1999; Hanski & Ovaskainen, 2003; Wiens, 1997). Transplant experiments are rarely used to test habitat quality at this scale, generally focusing on quality *within* putatively suitable sites; at this scale it is patchiness at the microhabitat level that could alter interpretation of transplant results. While not the scale traditionally considered in range-limit theory, there is observational evidence that declining microhabitat amount can impose range limits (Gillies et al., 2025). Further, for sessile and poorly-dispersing organisms (including *R. minor* and many other plants), the amount of suitable microhabitat within sites can strongly influence population viability and size (e.g. Grainger & Gilbert, 2017), which in turn affect a population’s ability to colonize adjacent sites (Alderman et al., 2005). We argue that small-scale patchiness is an underexplored component of range-limit dynamics, and important to consider given its potential influence on interpretating transplant results (Fig. 1). Ultimately, researchers must decide which spatial scale(s) are most relevant for their study species.

While we argued that testing both habitat quality and amount need not require substantial additional research effort, some additional effort may be needed to accurately estimate both. First, if microsite quality is quite variable, estimating both quality and amount may require additional plots per site. If redoing our own study, we would double the number of putatively-suitable plots. While we were generally successful in identifying suitable microsites, *a priori* identification was imperfect (some putatively-suitable plots had λ<1 even in sites where mean λ was ≥1; Fig. S5). At low-fitness sites in particular, more putatively-suitable plots would increase our ability to estimate quality of the best-available habitat. Second, whereas gradients in habitat quality can be inferred from gradients in fitness components (emergence, survival, reproduction) or proxies (e.g. growth), determining whether a microsite is suitable requires assaying lifetime fitness or population growth rates. Thus, depending on the study system, explicitly quantifying both the best-available habitat and habitat amount may require additional effort to capture enough suitable microsites if habitat is patchy (more plots/site), and to estimate fitness in longer-lived organisms (e.g. by increasing monitoring duration, or by transplanting multiple life stages to model population growth rates).

There is also a potential philosophical debate in measuring habitat quality and amount. When plot suitability is defined based on transplant performance, declines in the amount of suitable microhabitat could arise simply from declines in mean habitat quality. In other words, if mean λ declines, more plots will have λ<1 without obvious change in the apparent patchiness of a site. If so, it could be argued that patchiness is simply a by-product of habitat quality, and not inherently important on its own (Brown 1984). We argue the reverse; that whenever fitness varies among microsites, as it ubiquitously does when assayed experimentally (Bontrager et al., 2021), microsites are inherently patchy and patchiness is biologically interesting. Any increase in the proportion of unsuitable microsites increases the patchiness experienced by plants (and other organisms with undirected dispersal), as more propagules will land in unsuitable microsites. This will change the benefits of dispersal, just as happens at the landscape scale (Hargreaves & Eckert, 2014; Travis et al., 1999). Nevertheless, independent assessments of microsite availability (e.g. availability of suitable host plants or micro-climates) could help clarify whether declines in suitable microsites are due to changing resource distribution vs. declining mean suitability.

### Rhinanthus range-limit dynamics

While our aim was to test theory about habitat amount, our results also reveal unexpected performance declines at *R. minor*’s high-elevation range edge. Five years of transplants from 2011–2016 found that high-edge populations on Mt. Allan were self-sustaining, with fitness equivalent to mid- and low-elevation sites (Hargreaves & Eckert 2019; Ensing & Eckert 2019). In 2022 however, fitness was below replacement rates and significantly lower than at mid and low elevations. Similarly, simulations parameterized with 2022 data consistently underestimated *R. minor*’s “true” range limit, failing to expand through the entire range-edge section of the landscape. Lower fitness in 2022 is unlikely due to differences in transplant design (i.e. inclusion of evenly-spaced plots), as many evenly-spaced plots contained suitable hosts or natural *R. minor* plants, and simulations parameterized exclusively with data from putatively-suitable plots still failed to colonize the entire edge section (Fig. S6). Rather, the decline seems to reflect a true deterioration of site quality, and is paralleled by declines in natural *R. minor* population sizes from >1000 to <25 plants (Hargreaves, 2014; Rahn & Hargreaves, unpublished data).

This decline contrasts the widespread prediction that climate warming will improve habitat at the high-elevations of species’ ranges and enable upward range shifts (Chen et al., 2011; Grabherr et al., 1994). Upward shifts of high-elevation range limits are among the most commonly documented responses to climate warming (Hickling et al., 2006; Lawlor et al., 2024; Parmesan & Yohe, 2003), with exceptions usually interpreted as cases where the range was limited by factors other than temperature. Results for *R. minor* are particularly surprising given experimental evidence that performance above its range is constrained by cold (Hargreaves & Eckert 2019). Our finding that warming has not conferred a demographic advantage at *R. minor*’s high-elevation range edge awaits explanation, but cautions that species may not always escape climate change via upward range shifts.

### Recommendations for future work

We propose three simple recommendations to maximize transplant experiments’ power to detect the influence of both habitat quality and amount. **1)** As argued above, deliberately placing some plots in putatively-suitable or optimal microsites, and some evenly/randomly agnostic to microsite quality, enables better inference than either approach alone (see Fig. 1). In practice, many transplant experiments probably already employ something close to this design, deliberately sampling a variety of microhabitats. Thus our next recommendations apply to all experiments, regardless of the microsite-sampling strategy. **2)** Explicitly record and report which plots are in which type of microsite (many studies do not report how plot locations were selected), and ensure that the ratio is consistent across sites. This will enable readers to better interpret results and, if studies include both putatively-suitable and evenly/randomly placed plots, to disentangle changes in quality of the best-available habitat, habitat amount, and overall site quality. **3)** Include raw, plot-level habitat-quality data in figures (in addition to means and variation), so readers can distinguish changes in maximum microsite quality and the proportion of suitable microsites from changes in mean habitat quality. Transplant experiments have deeply enriched our understanding of species’ range dynamics; these relatively small changes to design and reporting could increase the explanatory power of these labour-of-love experiments, and advance our understanding of how habitat amount contributes to species’ range limits.

## Acknowledgements

We thank Anika Anderson, Sarah Chaddock, Maggie Blondeau and Shannon Meadley-Dunphy, for help executing our transplant experiment, Jake Alexander for helpful comments on an earlier version of the manuscript, and staff at the Nakiska Ski Area and Alberta Parks for allowing us to conduct research on Mt Allan (Permit #22-297). This research was funded by a Natural Sciences and Engineering Research Council (NSERC) Discovery Grant (ALH), Canadian Foundation for Innovation grant (ALH), Alberta Conservation Association Biodiversity Grant (OJR), NSERC CGS-M and CGS-D scholarships (OJR), and Student Undergraduate Research Awards from the McGill Biology Department and Faculty of Science.

## Literature Cited

Alderman, J., McCollin, D., Hinsley, S. A., Bellamy, P. E., Picton, P., & Crockett, R. (2005). Modelling the Effects of Dispersal and Landscape Configuration on Population Distribution and Viability in Fragmented Habitat. Landscape Ecology, 20(7), 857–870. 10.1007/s10980-005-4135-5

Betts, M. G., Fahrig, L., Hadley, A. S., Halstead, K. E., Bowman, J., Robinson, W. D., Wiens, J. A., & Lindenmayer, D. B. (2014). A species-centered approach for uncovering generalities in organism responses to habitat loss and fragmentation. Ecography, 37(6), 517–527. 10.1111/ecog.00740

Bontrager, M., Usui, T., Lee-Yaw, J. A., Anstett, D. N., Branch, H. A., Hargreaves, A. L., Muir, C. D., & Angert, A. L. (2021). Adaptation across geographic ranges is consistent with strong selection in marginal climates and legacies of range expansion. Evolution, 75(6), 1316–1333. 10.1111/evo.14231

Brooks ME, Kristensen K, van Benthem KJ, Magnusson A, Berg CW, Nielsen A, Skaug HJ, Maechler M, Bolker BM (2017). “glmmTMB Balances Speed and Flexibility Among Packages for Zero-inflated Generalized Linear Mixed Modeling.” The R Journal, 9(2), 378–400. doi:10.32614/RJ-2017-066.

Brown, J. H. (1984). On the Relationship between Abundance and Distribution of Species. The American Naturalist, 124(2), 255–279. 10.1086/284267

Carter, R. N., & Prince, S. D. (1981). Epidemic models used to explain biogeographical distribution limits. Nature, 293(5834), 644–645. 10.1038/293644a0

Chen, I.-C., Hill, J. K., Ohlemüller, R., Roy, D. B., & Thomas, C. D. (2011). Rapid Range Shifts of Species Associated with High Levels of Climate Warming. Science, 333(6045), 1024–1026. 10.1126/science.1206432

Colwell, R. K., Brehm, G., Cardelús, C. L., Gilman, A. C., & Longino, J. T. (2008). Global Warming, Elevational Range Shifts, and Lowland Biotic Attrition in the Wet Tropics. Science, 322(5899), 258–261. 10.1126/science.1162547

Davis, M. B., & Shaw, R. G. (2001). Range Shifts and Adaptive Responses to Quaternary Climate Change. Science, 292(5517), 673–679. 10.1126/science.292.5517.673

Davison, P. J., & Field, J. (2018). Environmental barriers to sociality in an obligate eusocial sweat bee. Insectes Sociaux, 65(4), 549–559. 10.1007/s00040-018-0642-7

Early, R., & Sax, D. F. (2014). Climatic niche shifts between species’ native and naturalized ranges raise concern for ecological forecasts during invasions and climate change. Global Ecology and Biogeography, 23(12), 1356–1365. 10.1111/geb.12208

Ensing, D. J., & Eckert, C. G. (2019). Interannual variation in season length is linked to strong co-gradient plasticity of phenology in a montane annual plant. New Phytologist, 224(3), 1184–1200. 10.1111/nph.16009

Ensing, D. J., Sora, D. M. D. H., & Eckert, C. G. (2021). Chronic selection for early reproductive phenology in an annual plant across a steep, elevational gradient of growing season length. Evolution 75(7), 1681–1698. 10.1111/evo.14274

Falk, L. 2013. The Intensity and Effects of Insect Leaf Herbivory on Rhinanthus minor along its Elevational Range. [Honours thesis, Queens University]

Fox J., Weisberg S (2019). An R Companion to Applied Regression, Third edition. Sage, Thousand Oaks CA.

Gibson, C. C., & Watkinson, A. R. (1989). The host range and selectivity of a parasitic plant: Rhinanthus minor L. Oecologia, 78(3), 401–406. 10.1007/BF00379116

Gillies, G. J., Dungey, M. P., & Eckert, C. G. (2025). Evidence That Metapopulation Dynamics Maintain a Species’ Range Limit. Ecology Letters, 28(5), e70128. 10.1111/ele.70128

Grabherr, G., Gottfried, M., & Pauli, H. (1994). Climate effects on mountain plants. Nature, 369(6480), 448–448. 10.1038/369448a0

Grainger, T. N., & Gilbert, B. (2017). Multi scale responses to warming in an experimental insect metacommunity. Global Change Biology, 23(12), 5151–5163. 10.1111/gcb.13777

Griggs, R. F. 1914. Observations on the behavior of some species at the edges of their ranges. Bull. Torrey Bot. Club 41, 25–49

Grinnell, J. (1917). The Niche-Relationships of the California Thrasher. The Auk, 34(4), 427–433. 10.2307/4072271

Hampe, A., & Petit, R. J. (2005). Conserving biodiversity under climate change: The rear edge matters. Ecology Letters, 8(5), 461–467. 10.1111/j.1461-0248.2005.00739.x

Hanski, I. (1999). Metapopulation Ecology. OUP Oxford.

Hanski, I., & Ovaskainen, O. (2003). Metapopulation theory for fragmented landscapes. Theoretical Population Biology, 64(1), 119–127. 10.1016/S0040-5809(03)00022-4

Hargreaves, A. L. (2014). Evolutionary ecology of range limits: Conceptual syntheses and empirical tests. [PhD thesis, Queens University]

Hargreaves, A. L., Bailey, S. F., & Laird, R. A. (2015). Fitness declines towards range limits and local adaptation to climate affect dispersal evolution during climate induced range shifts. Journal of Evolutionary Biology, 28(8), 1489–1501. 10.1111/jeb.12669

Hargreaves, A. L., & Eckert, C. G. (2014). Evolution of dispersal and mating systems along geographic gradients: Implications for shifting ranges. Functional Ecology, 28(1), 5–21. 10.1111/1365-2435.12170

Hargreaves, A. L., & Eckert, C. G. (2019). Local adaptation primes cold-edge populations for range expansion but not warming-induced range shifts. Ecology Letters, 22(1), 78–88. 10.1111/ELE.13169

Hargreaves, A. L., Samis, K. E., & Eckert, C. G. (2014). Are species’ range limits simply niche limits writ large? A review of transplant experiments beyond the range. American Naturalist, 183(2), 157–173. 10.1086/674525

Hargreaves, A. L., Weiner, J. L., & Eckert, C. G. (2015). High-elevation range limit of an annual herb is neither caused nor reinforced by declining pollinator service. Journal of Ecology, 103(3), 572–584. 10.1111/1365-2745.12377

Hickling, R., Roy, D. B., Hill, J. K., Fox, R., & Thomas, C. D. (2006). The distributions of a wide range of taxonomic groups are expanding polewards. Global Change Biology, 12(3), 450–455. 10.1111/j.1365-2486.2006.01116.x

Holt, R. D., Barfield, M., & Gonzalez, A. (2003). Impacts of environmental variability in open populations and communities: “Inflation” in sink environments. *Theoretical Population Biology*, Understanding the Role of Environmental Vatiation in Population and Community Dynamics, 64(3), 315–330. 10.1016/S0040-5809(03)00087-X

Holt, R. D., & Keitt, T. H. (2000). Alternative causes for range limits: A metapopulation perspective. Ecology Letters, 3(1), 41–47. 10.1046/J.1461-0248.2000.00116.X/FORMAT/PDF

Hulst, R. V., Thériault, A., & Shipley, B. (1986). The systematic position of the genus *Rhinanthus* (Scrophulariaceae) in North America. Canadian Journal of Botany, 64(7), 1443–1449. 10.1139/b86-196

Humboldt, A.v. & Bonpland, A. (1805). Essai sur la géographie des plantes: accompagné d’un tableau physique des régions équinoxiales, fondé sur des mesures exécutées, depuis le dixième degré de latitude boréale jusqu’au dixième degré de latitude australe, pendant les années 1799, 1800, 1801, 1802 et 1803. Levrault, Schoell, and Company, Paris.

Lajoie, G., & Vellend, M. (2018). Characterizing the contribution of plasticity and genetic differentiation to community-level trait responses to environmental change. Ecology and Evolution, 8(8), 3895–3907. 10.1002/ece3.3947

Lawlor, J. A., Comte, L., Grenouillet, G., Lenoir, J., Baecher, J. A., Bandara et al. (2024). Mechanisms, detection and impacts of species redistributions under climate change. Nature Reviews Earth & Environment, 5(5), 351–368. 10.1038/s43017-024-00527-z

Lechowicz, M. J., & Bell, G. (1991). The Ecology and Genetics of Fitness in Forest Plants. II. Microspatial Heterogeneity of the Edaphic Environment. Journal of Ecology, 79(3), 687–696. 10.2307/2260661

Lee-Yaw, J. A., Kharouba, H. M., Bontrager, M., Mahony, C., Csergő, A. M., Noreen et al. (2016). A synthesis of transplant experiments and ecological niche models suggests that range limits are often niche limits. Ecology Letters, 19(6), 710–722. 10.1111/ele.12604

Lennon, J. J., Turner, J. R. G., & Connell, D. (1997). A Metapopulation Model of Species Boundaries. Oikos, 78(3), 486. 10.2307/3545610

Lenth, R., Piaskowski J (2025). emmeans: Estimated Marginal Means, aka Least-Squares Means. doi:10.32614/CRAN.package.emmeans, R package version 2.0.1.

Levin, D. A., & Schmidt, K. P. (1985). Dynamics of a Hybrid Zone in *Phlox*: An Experimental Demographic Investigation. American Journal of Botany, 72(9), 1404–1409. 10.2307/2443513

Lüdecke, et al., (2021). performance: An R Package for Assessment, Comparison and Testing of Statistical Models. Journal of Open Source Software, 6(60), 3139. 10.21105/joss.03139

McLane, S. C., & Aitken, S. N. (2012). Whitebark pine () assisted migration potential: Testing establishment north of the species range. Ecological Applications, 22(1), 142–153. 10.1890/11-0329.1

Oldfather, M. F., Van Den Elzen, C. L., Heffernan, P. M., & Emery, N. C. (2021). Dispersal evolution in temporally variable environments: Implications for plant range dynamics. American Journal of Botany, 108(9), 1584–1594. 10.1002/ajb2.1739

Paquette, A., & Hargreaves, A. L. (2021). Biotic interactions are more often important at species’ warm versus cool range edges. Ecology Letters, 24(11), 2427–2438. 10.1111/ele.13864

Parmesan, C., & Yohe, G. (2003). A globally coherent fingerprint of climate change impacts across natural systems. Nature, 421(6918), Article 6918. 10.1038/nature01286

Persi, J. (2020). Elevational patterns in seed fates: experimental test in the Rocky Mountains [MSc Thesis, McGill University]

Phillips, B. L. (2012). Range shift promotes the formation of stable range edges. Journal of Biogeography, 39(1), 153–161. 10.1111/j.1365-2699.2011.02597.x

Pironon, S., Papuga, G., Villellas, J., Angert, A. L., García, M. B., & Thompson, J. D. (2017). Geographic variation in genetic and demographic performance: New insights from an old biogeographical paradigm. Biological Reviews, 92(4), 1877–1909. 10.1111/brv.12313

Rapoport, E.H. (1982) Aerography: Geographical Strategies of Species. Pergamon Press, New York.

R Core Team. (2025) R: A Language and Environment for Statistical Computing. R Foundation for Statistical Computing.

Samis, K. E., & Eckert, C. G. (2009). Ecological correlates of fitness across the northern geographic range limit of a Pacific coast dune plant. Ecology, 90(11), 3051–3061. 10.1890/08-1914.1

Saona, N. M. (2002). Host selection of the hemiparasitic plant, Rhinanthus minor. [MSc Thesis, University of Calgary]

Soberón, J. (2007). Grinnellian and Eltonian niches and geographic distributions of species. Ecology Letters, 10(12), 1115–1123. 10.1111/j.1461-0248.2007.01107.x

Solarik, K. A., Messier, C., Ouimet, R., Bergeron, Y., & Gravel, D. (2018). Local adaptation of trees at the range margins impacts range shifts in the face of climate change. Global Ecology and Biogeography, 27(12), 1507–1519. 10.1111/geb.12829

Tito, R., Vasconcelos, H. L., & Feeley, K. J. (2021). Multi-population seedling and soil transplants show possible responses of a common tropical montane tree species (*Weinmannia bangii*) to climate change. Journal of Ecology, 109(1), 62–73. 10.1111/1365-2745.13443

Travis, J. M. J., Murrell, D. J., & Dytham, C. (1999). The evolution of density–dependent dispersal. Proceedings of the Royal Society B: Biological Sciences, 266(1431), 1837–1842. 10.1098/rspb.1999.0854

Venables, W. N., & Ripley, B. D. (2002). Modern Applied Statistics with S. Springer. 10.1007/978-0-387-21706-2

Westbury, D. B. (2004). *Rhinanthus minor* L. Journal of Ecology, 92(5), 906–927. 10.1111/j.0022-0477.2004.00929.x

Whittaker, R. H. (1956). Vegetation of the Great Smoky Mountains. Ecological Monographs, 26(1), 1–80. 10.2307/1943577

Wiens, J. A. (1976). Population Responses to Patchy Environments. Annual Review of Ecology and Systematics 7, 81–120.

Wiens, J. A. (1997). Metapopulation Dynamics and Landscape Ecology. In: Metapopulation Biology. {eds Hanski, I., Gilpin M}. Elsevier, pp. (43–61)

Zeileis, A., Hothorn, T. (2002). Diagnostic Checking in Regression Relationships. R News 2(3), 7–10. URL https://CRAN.R-project.org/doc/Rnews/

