## Supplemental Material for "A modified transplant design reveals that habitat quality and quantity both limit a species’ range"

### Section 1: Supplementary tables

**Table S1: Statistical results for the effect of seed source on performance.**

Transplants might detect low fitness even at high-quality sites if plants at those sites produced poor quality seeds in the preceding year. To disentangle the effects of site quality in 2022 from the quality of seeds produced in 2021, we planted both local seeds (20/plot) and standard seeds (5/plot, all from Mid1) in each plot at each site. We tested whether the detected pattern of fitness across sites varied depending on seed source (local vs. standard) using negative binomial GLMs. The full model was *seeds produced per plot ~ site x seed source + (1|plot)*, with an offset for the log(number of seeds planted per source). We compared models with and without the terms of interest using likelihood ratio tests. Neither the interaction term nor seed source significant improved model fit, thus local and standard seeds detected similar fitness patterns among sites and standard seeds had similar overall performance to local seeds. All other analyses use local seeds only.

| Model fixed effects | Chi^2^ | P |
| --- | --- | --- |
| M1) seeds produced ~ site + seed source + (1\|plot)*, df=10* | | |
| M2) seeds produced ~ site x seed source + (1\|plot)*, df = 18* | | |
| M3) seeds produced ~ site + (1\|plot)*, df=10* | | |
| M1 vs. M2: Interaction term | 12.82 | 0.12 |
| M2 vs. M3: seed source | 12.819 | 0.1182 |

**Table S2: Simulation model results from final generation for each simulation model type.** Numbers in brackets represent measures of standard deviation. The final generation was generation 1200 for simulations presented in the main results (quality only, amount only, and quality + amount) but 600 for supplementary simulations (combined sampling vs putatively suitable sampling, described in Supplementary Methods 2 and Figure S6)

| Landscape type | Generation | Mean cells occupied beyond row 300 | Mean furthest row reached | Mean cells occupied across landscape | | Mean density of occupied cells across landscape |
| --- | --- | --- | --- | --- | --- | --- |
| **Testing gradients in habitat quality vs. amount** | | |  |  |  | |
| Quality only | 1200 | 0 | 206.825  (± 1.8) | 5111.2  (± 20.4) | | 27.8  (± 0.6) |
| Amount only | 1200 | 0 | 215.425  (± 6.9) | 4869.2  (± 18.1) | | 43.4  (± 0.7) |
| Quality + amount | 1200 | 0 | 202.9875  (± 100.0) | 4727.0  (± 19.5) | | 29.6  (± 0.5) |
| Quality only | 100 | 0 | 164.100  (± 6.7) | 3842.0  (±120.2) | | 32.7  (± 0.9) |
| Amount only | 100 | 0 | 196.225  (± 5.1) | 4441.0  (± 124.8) | | 43.0  (± 0.7) |
| Quality and amount | 100 | 0 | 153.900  (± 5.8) | 3433.450  (± 100.0) | | 35.811  (± 0.9) |
| **Testing effect of transplant design** | | |  |  |  | |
| “Combined” sampling | 600 | 0 | 203.925  (± 1.3) | 5647.75  (± 72.216) | | 21.88022  (± 0.4615082) |
| “Putatively-suitable” sampling | 600 | 0 | 204.45  (± 1.1) | 8749.5  (± 78.9) | | 29.14233  (± 0.7) |

### Section 2: Supplementary figures

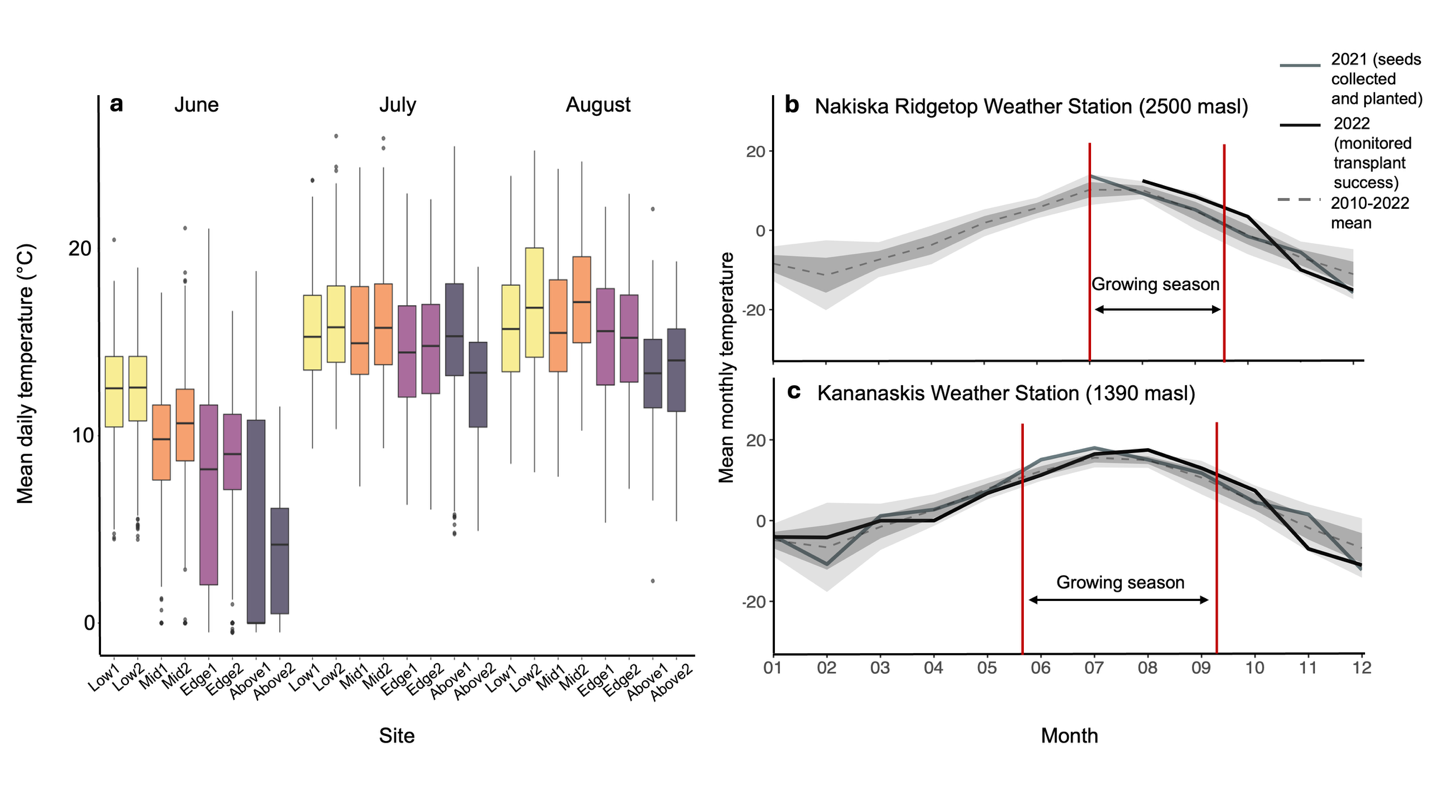

**Figure S1: Local temperatures during study period. (a)** Mean temperatures recorded by iButton temperature loggers at transplant sites during *R. minor*’s growing season. iButtons recorded air temperature close to the ground surface every 2 hours during summer 2022 at 10 to 20 plots per site depending on logger availability (n=6 iButtons at Above2 as many were faulty). iButton loggers were covered in white electrical tape to reduce solar reflection, taped horizontally on top of a metal nail, and the nail was pushed into the ground such that the iButton was ~1 cm above the ground surface. Thick black lines, boxes, and whiskers represent means, 1^st^ and 3^rd^ quartiles, and upper and lower range of the data excluding potential outliers, respectively. (**b**–**c)** Temperature trends at two weather stations close to our transplant sites in 2021 (year of seed collection and planting) and 2022 (year of transplant monitoring); data downloaded from Environment and Climate Change Canada, 2023. To determine whether either year was anomalous, we compared the monthly average temperatures recorded at each station to that station’s decadal averages. We calculated monthly averages by taking the average of all daily averages (the grand mean) across each month. The grand mean is shown as a solid black line for individual study years and a dashed line for the 2010-2022 mean. Dark and light grey shading show 1 and 2 standard deviations, respectively. Our monitoring year (2022) was warm compared to the past decade, especially at high elevations and during July and August. ECCC data at Nakiska Ridgetop were not available from January-June 2021 (inclusive) or January-July 2022 (inclusive).

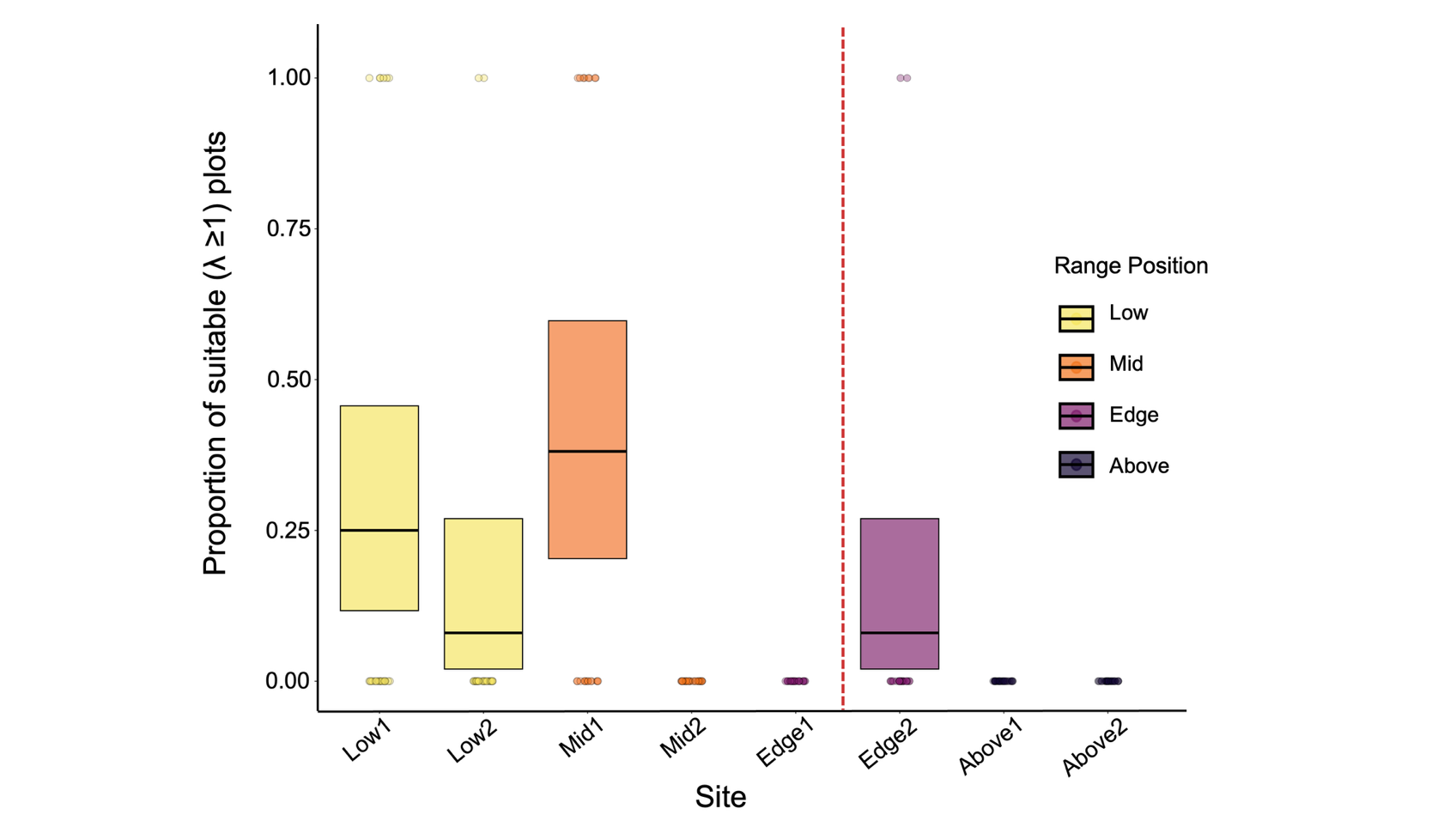

**Figure S2: Habitat amount detected by evenly-spaced plots only.** Black bars and boxes show estimated marginal means and 95% CI of a binomial model, which considered only sites that had at least one evenly-spaced plot with positive population growth (i.e. λ ≥1). Points show raw data for all sites. The probability of a plot being suitable (i.e. λ ≥1) varied among sites (χ^2^_df = 3_ = 9.46, p = 0.024), but no pairwise contrasts were significant. Trends were similar if models included all plots (putatively-suitable and evenly-sampled; Figure 3c).

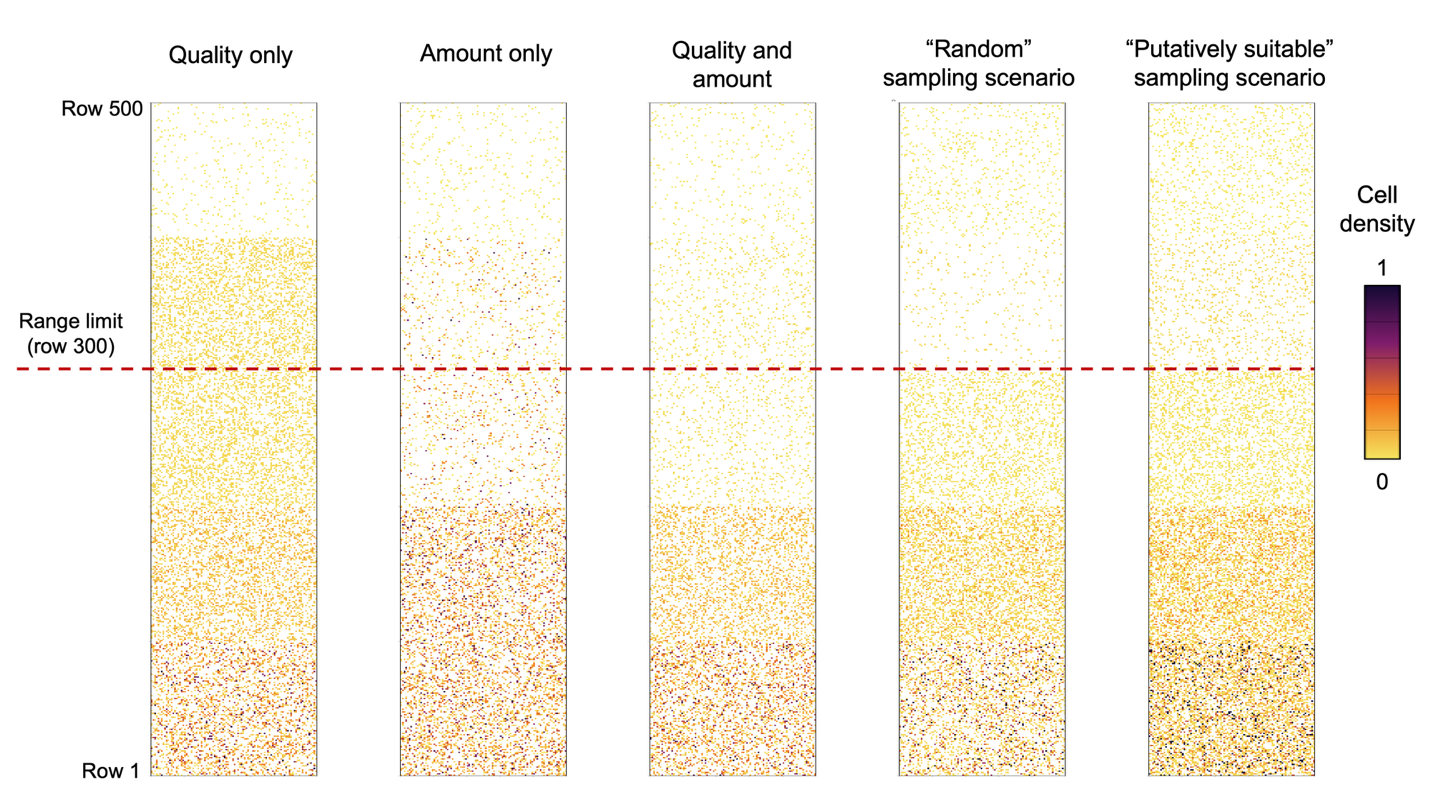

**Figure S3:** **Example model landscapes for each simulation type**. Each landscape is made up of 5 sections corresponding to *R. minor*’s Low, Mid, Edge, Above1 and Above2 range sections. The leftmost three landscapes compare expansion across landscapes where range sections differ in the quality of suitable cells, the amount of suitable cells, or both (main analyses; Fig. 5). The rightmost two landscapes represent the landscapes one would infer from a transplant experiment using our full design (putatively-suitable + evenly-spaced) vs. an experiment using only putatively-suitable plots, and are used to test the effect of transplant design (Figure S7, Supplementary Methods 2). Colour scale in legend shows the relative quality of each cell; values <0.001 are masked to white to improve visualization. Dashed red line shows the actual location of *R. minor*’s high-elevation range limit (i.e. between Edge and Above1 range sections).

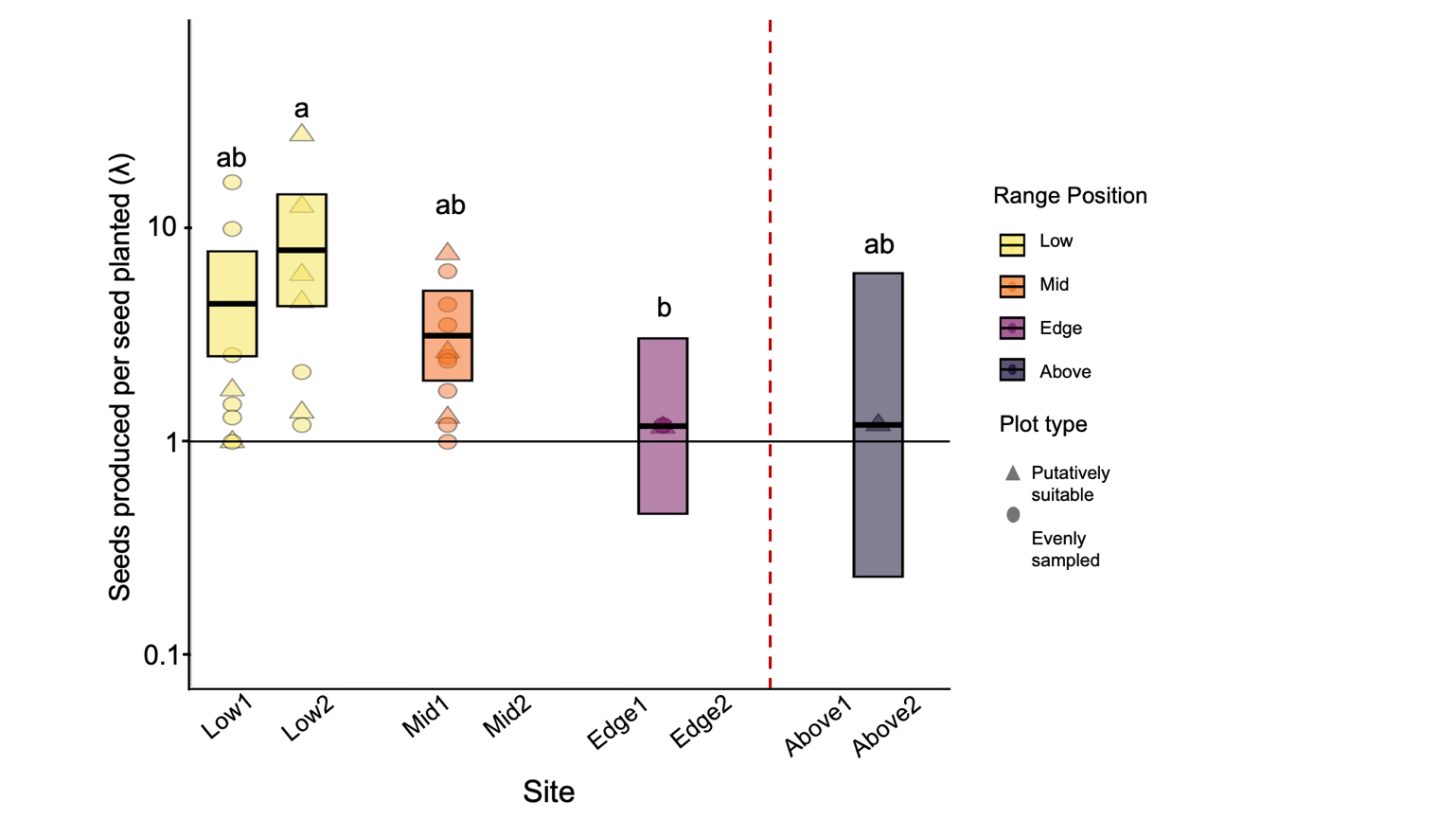

**Figure S4: Habitat quality in plots with λ ≥1.** To estimate the proportion of suitable plots in each site, we had to define ‘suitable’. Here, suitable plots are defined as those where *R. minor* transplant fitness λ ≥1, whether or not plots were selected as putatively-suitable or evenly-spaced (Fig. 3b shows results when ‘suitable’ is defined as plots where *R. minor* transplants produced fruits). Black bars and boxes show estimated marginal means and 95% confidence intervals from a negative binomial GLM; points show raw data for each plot. Horizontal reference line at λ=1 differentiates plots in which population growth was self-sustaining (λ≥1) or not (λ<1). Confidence intervals are particularly large at range-edge and above-range sites as these sites had few plots with λ ≥1 and therefore small sample sizes for this analysis. Dashed red line indicates *R. minor*’s high-elevation range limit on Mt. Allan.

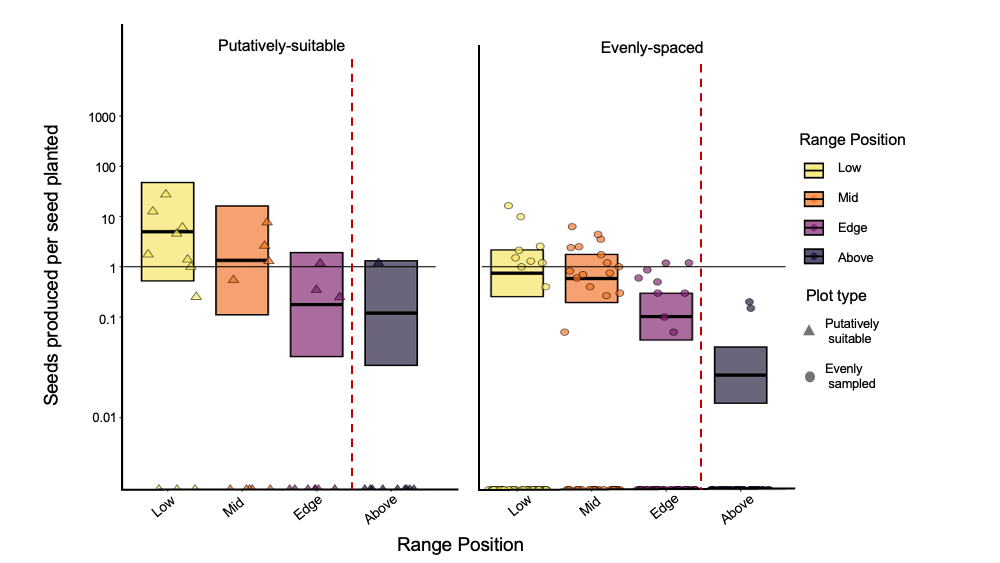

**Figure S5: Range-wide habitat quality detected by putatively-suitable vs. evenly-spaced transplant plots.** At each site we planted 5 plots into microhabitats identified *a priori* as putatively suitable for *R. minor*, based on the presence of naturally-occurring reproductive *R. minor* and/or suitable host plants, and 25 plots evenly spaced across the site, agnostic to microhabitat quality. Including plot type as a predictor improved model fit (likelihood ratio test: χ^2^_df=1_ = 7.37, *P* = 0.0066), and mean fitness was higher in putatively-suitable plots (*plot type*: χ^2^_df=1_ = 5.94, p = 0.015), confirming that we were generally able to identify suitable (or at least the best available) habitat for *R. minor*. However, some plots identified as putatively-suitable had λ<1, even at sites that with mean λ>1 (e.g. sites in the ‘Low’ and ‘Mid’ range positions), indicating that this ability was imperfect. Qualitatively, evenly-spaced suitable plots detected a slightly steeper fitness decline from range edge to beyond the range, but the *range position x plot type* interaction was not significant (likelihood ratio test: χ^2^_df=3_ 1.71, *P* = 0.634). Thick black lines and shaded rectangles show the estimated marginal means and 95% confidence intervals back-transformed from a negative binomial GLM. Points show raw λ data for each plot. Horizontal reference line shows λ=1, below which populations are not self-sustaining (i.e. habitat unsuitable).

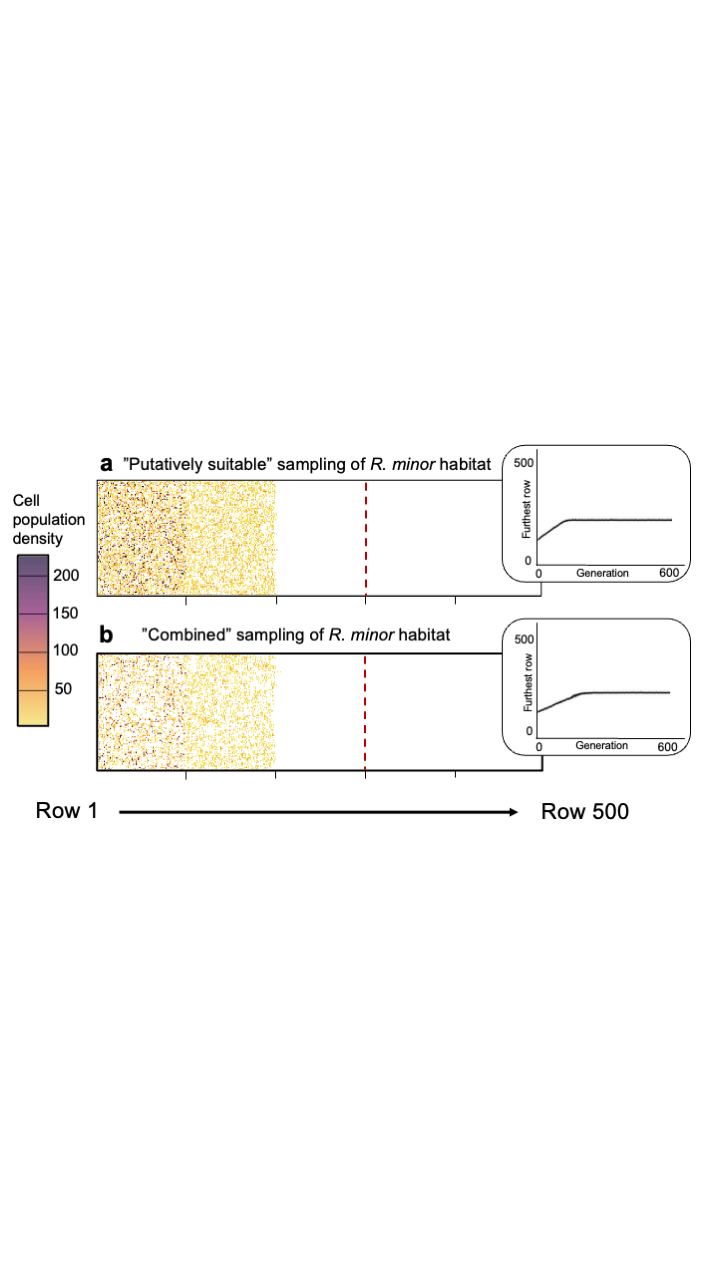

**Figure S6: Simulation results exploring the effect of transplant design.** Simulated *R. minor* range expansion across landscapes parameterized with field data from either **(a)** only putatively suitable plots or **(b)** putatively-suitable and evenly-spaced plots. As in other simulation models (Fig. 5), landscapes are divided into five equal range sections (Low, Mid, Edge, Above1, Above2). Landscape plots show densities in all cells at generation 600 for one example model run. Dashed red lines show *R. minor*’s “true” range limit at row 300, i.e. between the Edge and Above-range sections. Insets: accumulation curves showing the furthest row of the landscape reached at each generation in each scenario (mean ± 2 standard deviations across 10 randomly selected runs per landscape type). The location of the range limit at generation 600 was similar for both scenarios, but simulated expansion across landscapes parameterized with only putatively suitable plots was slightly faster (see insets) and the final predicted population size and density in occupied cells was higher than for landscapes parameterized with data from all plots (see Table S2).

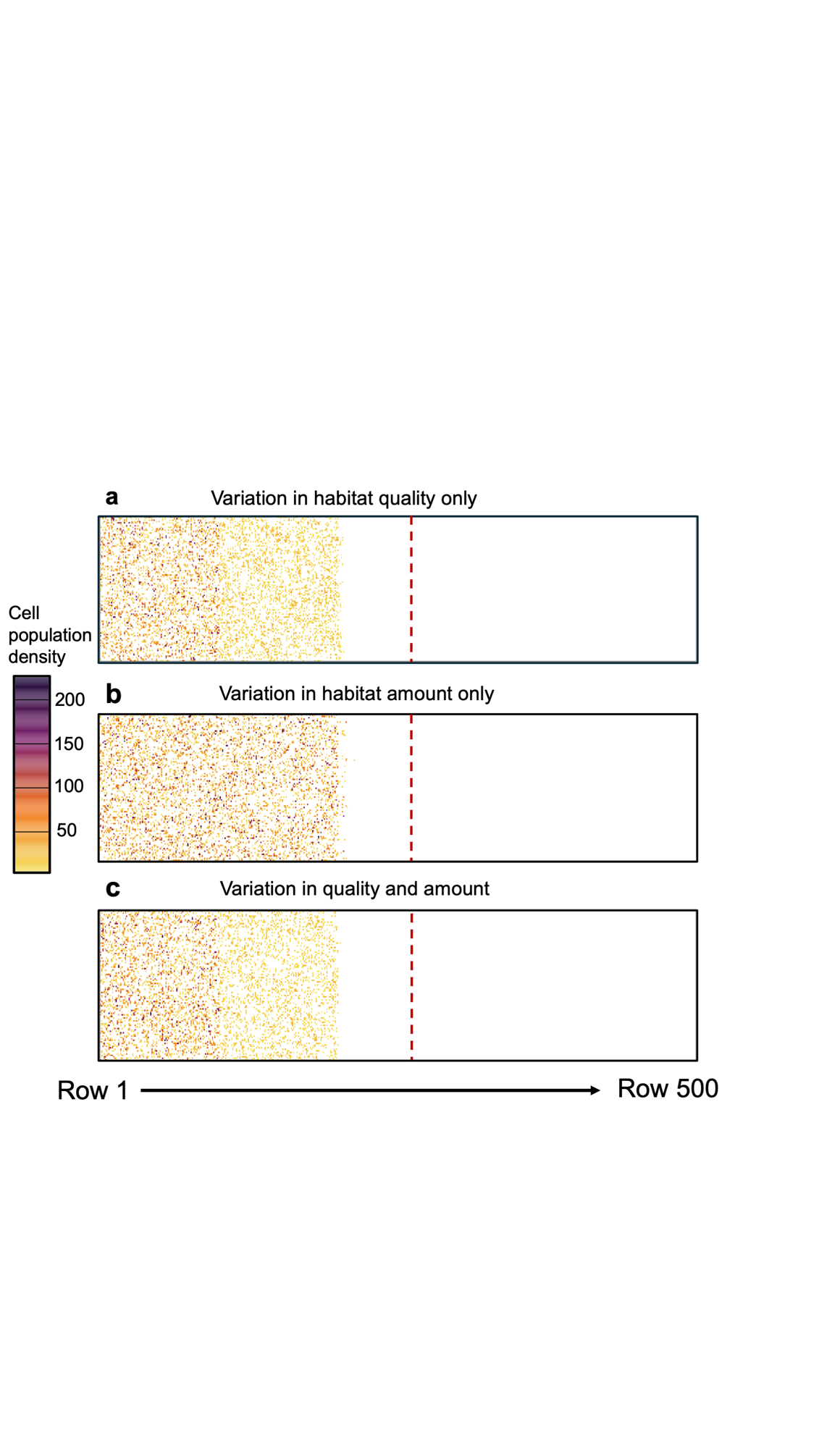

**Figure S7:** **Simulated *R. minor* range expansion at generation 1200.** The arrangement of landscapes is the same as in Figure 5 (which shows cell densities at generation 100), but these plots show cell densities at the final generation of each run (generation 1200; statistical results in Table S3). Landscape-wide gradients in habitat quality alone and amount alone both produced stable range limits by generation 1200 of simulated range expansion.

### Section 3: Supplementary Text

### Methods S1) Details about simulation models

#### Simulation model details

We ran 40 replicates of each landscape type, which optimized use of the computer cluster available to us (the Digital Research Alliance Canada Narval Cluster) when accounting for memory needs and computation time. To start each simulation, we randomly seeded 1000 individuals across the first 100 rows of their respective landscape (i.e. the Low section). While other expansion models have used an initial generation that involves most cells in the landscape being at carrying capacity, starting with very large numbers of individuals quickly exceeded our computational power. We often find very small edge populations of *R. minor,* and therefore a smaller starting population size representing an expanding front seemed more realistic.

Each seeded individual began as a mature plant, which then reproduced. The number of offspring was randomly drawn from a Poisson distribution with a mean calculated using a modified Hassel-Commins population growth equation (see main text). In this equation, we scaled fixed values of R_max_ and K by the habitat quality value of the cell, allowing us to incorporate the effect of habitat quality on individual reproduction. We initially used fixed values of K=521 and R_max_=225, which represent the highest observed values for reproductive individuals per 1 m^2^ radius and seeds produced per plant during previous work with *R. minor* in our study area. However, these resulted in excessive computation times, so we reduced values by a factor of 2.25 (to K=230 and R_max_=100). The Hassel-Commins equation incorporates negative density dependence, such that mean fitness declines as cell density increases. This also introduces priority effects, as initial dispersers into the cell (i.e. that establish at low densities) are more likely to be assigned high fitness. To incorporate seed death over winter (e.g. due to post-dispersal seed predation), only 50% of offspring were allowed to survive (selected randomly for each plant). Surviving offspring then dispersed in a random direction (up, down, left or right). Dispersal distance was drawn from a Poisson distribution with mean=1, such that most seeds land in the same cell as their parent, rare long-distance dispersal is possible (which matches presumed dispersal dynamics for *R. minor*; Westbury, 2004). If individuals dispersed beyond the lower or upper limits of the landscape, they were returned to their closest row to reproduce. If individuals dispersed beyond the left or right boundaries of the landscape, they simply wrapped to the other side (Hargreaves et al., 2015). Individuals were added to the cells they dispersed into so long as the cell was not already at carrying capacity. Individuals reproduced in their new cell, and the simulation continued in this manner for 1200 generations. We selected parameter values and model behaviour to approximate real-world *R. minor* biology as closely as possible, given limits on computing time and power; full exploration of the model’s parameter space was beyond the scope of this study.

#### Accounting for instability in the modified Hassel-Commins equation

Specific combinations of population density and cell quality resulted in instability within the Hassel-Commins equation or biologically unrealistic results. Generally, this occurred if low cell quality (<0.01, equivalent to a transplant plot that produced less than 1 seed per seed planted), caused the denominator of the equation to approach 0. Denominator values of the HC equation very close to 0 resulted in -INF or +INF. Negative λ estimates also arose if a cell’s quality was < 0.01 and it already had high density – this result was not valid in our simulation as the Poisson draw we did to simulate offspring production during the simulation requires positive values. Finally, cell quality values of exactly 0.01 would produce fitness values of 1 regardless of density, and values that were very close to but slightly below 0.01 resulted in fitness values that were much larger than biologically realistic. We address these issues in two ways. First, as +INF, -INF or negative values all resulted from a combination of extremely low cell quality and high cell density, both conditions that promote extremely low fitness, we replaced these values with 0. Second, to ensure no large, biologically unrealistic values were being produced as a result of cell quality values that were very close to but < 0.01, we replaced cell values between 0.009 and 0.01 (inclusive) with 0.0101. This adjustment still reflects extremely low cell quality, but avoids artificial inflation of fitness.

### Methods S2) Using simulations to explore the impact of sampling design.

A secondary goal for our simulation was to test whether a transplant design that explicitly quantified habitat amount (i.e. incorporated evenly-spaced plots) would yield different conclusions about *R. minor*’s range limit than a design that used only putatively-suitable microsites. To do so, we simulated range expansion across landscapes that represented the results of the two sampling designs. Design 1 represents a landscape that researchers would infer from results of putatively-suitable plots. We drew quality values (λ from transplant plots, as described in main methods text) for each range section from only putatively suitable plots in that section. To increase sample size and variation, we used values from both the 5 putatively suitable plots identified *a priori* plus the five putatively best-quality evenly-spaced plots, selected based on pre-planting observations of natural *R. minor* and/or appropriate hosts recorded during transplant setup (i.e. blind to actual transplant performance). Design 2 represents a landscape inferred from our modified transplant design, including both putatively-suitable and evenly-spaced plots. For this landscape, we drew λ values from all transplant plots within each range section.

As done for the landscapes presented in the main text, we scaled all lifetime fitness values to be between 0 and 1 and created 40 replicates of each landscape type. We then used the same simulation described in the main text and in Methods S1 to identify differences in simulated range expansion between the two landscape designs. We stopped the simulation at generation 600, but expansion for both landscape types (Design 1 and Design 2) plateaued between generations 200-250.

#### Results

The extent of range expansion at generation 600 was similar between Design 1 & 2 (Figure S7). However, Design 1 (putatively-suitable microsites only) had higher total population size and density of occupied cells (Table S2) and faster range expansion (see insets in Figure S7). This suggests that incorporating evenly-spaced plots yields more conservative estimates of (a) overall site quality and (b) the speed of future range expansion.

**Literature cited**

Data from: Environment and Climate Change Canada (2023). Available at [https://climate.weather.gc.ca/](https://climate.weather.gc.ca/historical_data/search_historic_data_e.html)

Hargreaves, A. L., Bailey, S. F., & Laird, R. A. (2015). Fitness declines towards range limits and local adaptation to climate affect dispersal evolution during climate‐induced range shifts. *Journal of Evolutionary Biology*, *28*(8), 1489–1501. https://doi.org/10.1111/jeb.12669

Lenth, R., Piaskowski J (2025). emmeans: Estimated Marginal Means, aka Least-Squares Means. doi:10.32614/CRAN.package.emmeans, R package version 2.0.1, <https://CRAN.R-project.org/package=emmeans>

Westbury, D. B. (2004). Rhinanthus minor L. *Journal of Ecology*, *92*(5), 906–927. https://doi.org/10.1111/j.0022-0477.2004.00929.x
